# A novel *Pbunavirus* phage from the Ganges River in biocontrol of multidrug-resistant *Pseudomonas aeruginosa* in wastewater by downregulation of *las/rhl* systems and reduction of KPC carbapenemases

**DOI:** 10.1101/2024.11.01.621487

**Authors:** Rachel Samson, Sai Joshi, Rinka Pramanik, Krishna Khairnar, Mahesh Dharne

## Abstract

Wastewater systems are reservoirs of multidrug-resistant (MDR) pathogens, including *Pseudomonas aeruginosa*, mirroring those found in the hospital effluents. Effective treatment of raw sewage for MDR *P. aeruginosa* abatement is critical for safe effluent disposal. This study reports the isolation of a novel phage (ɸPRS-1) within the *Pbunavirus* genus from a pristine stretch of the Ganges River, targeting the MDR strain of *P. aeruginosa*. Antibacterial and antibiofilm assays of ɸPRS-1 against *P. aeruginosa* PAO1 demonstrated robust efficacy, achieving log_10_ reductions of 8.14 in planktonic cells, 9.83 in biofilm inhibition, and 7.01 in biofilm disruption after 24 hours. Additionally, ɸPRS-1 effectively disrupted the preformed biofilm of various other MDR *P. aeruginosa* strains with >5 log_10_ reductions. Mechanistically, ɸPRS-1 downregulated quorum-sensing genes and reduced the carbapenemase resistance KPC cluster, validated through qRT-PCR analysis. Evaluation of ɸPRS-1 in raw sewage revealed a 4.2 log_10_ CFU/mL reduction in the *P. aeruginosa* counts, highlighting its efficacy in complex environmental matrices. *In vitro,* cytotoxicity assays confirmed the phage’s safety. The use of ɸPRS-1 for tackling MDR *P. aeruginosa* in the wastewater offers a sustainable alternative, potentially reducing reliance on other treatment methods. The findings have attempted to deliver SDGs 3, 6, 11, 14, and 15.

**Highlights:**

- A novel phage (ɸPRS-1) from an untapped location of the Ganges River
- ɸPRS-1 reduces planktonic cells of MDR *P. aeruginosa* by 8.14 log_10_ in 24 hours
- Antibiofilm potentials: 9.83 log_10_ reduction (inhibition) >5 log_10_ reduction (disruption)
- Downregulates quorum-sensing genes and reduces KPC resistance in MDR *P. aeruginosa*
- ɸPRS-1 shows 4.2 log_10_ CFU/mL reduction of MDR *P. aeruginosa* in raw sewage

**Graphical Abstract:** 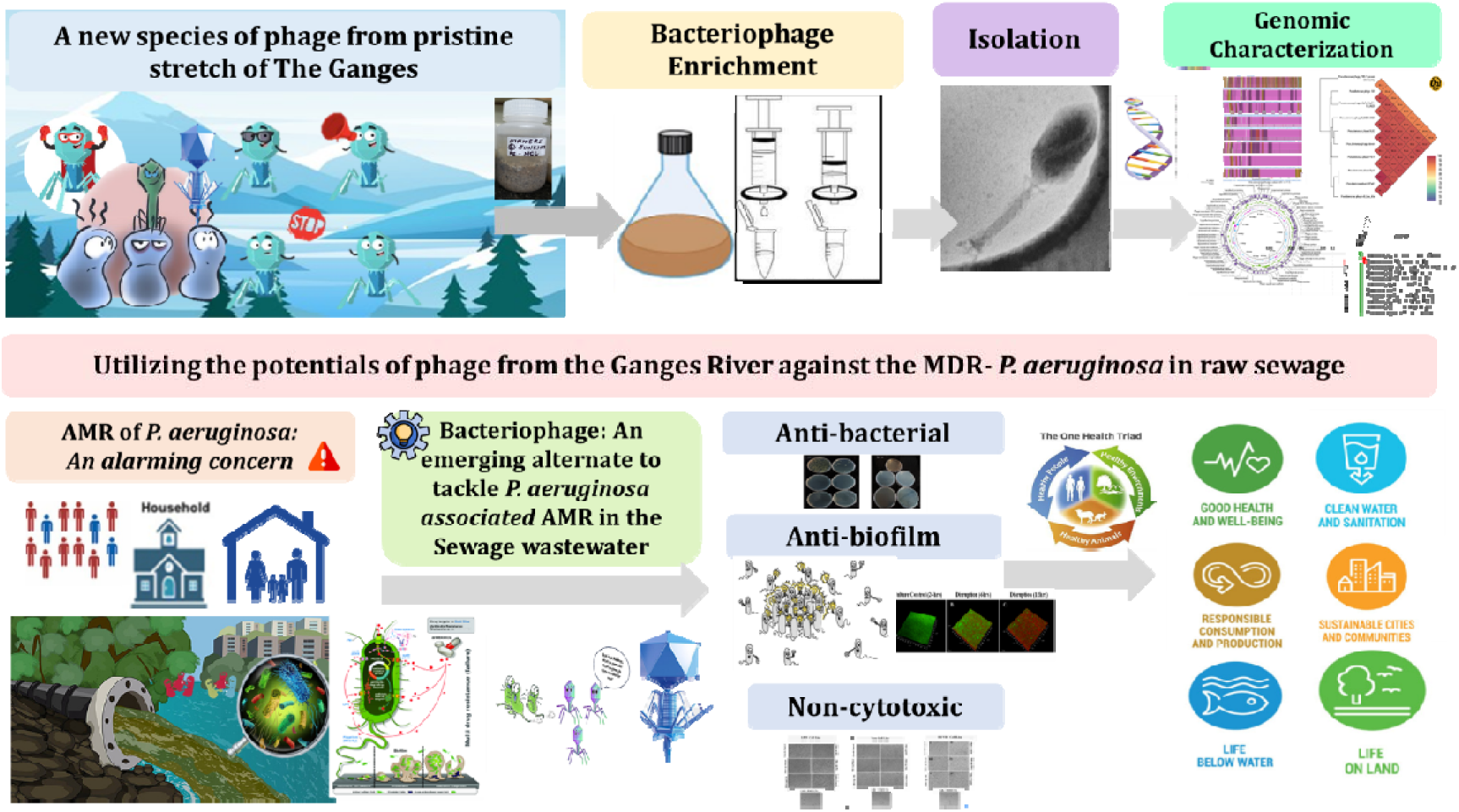

## 1. Introduction

Antimicrobial resistance (AMR) ranks among the foremost global public health concerns by the World Health Organisation (WHO), posing a significant barrier to attaining universal healthcare and sustainable development goals (SDGs) like SDGs 2,3,6,8, 11,12, 14, and 15 [1]. Human activities primarily influence aquatic ecosystems, serving as hotspots for the proliferation and evolution of antimicrobial resistance genes (ARGs) and associated bacteria. Water security is a key component of environmental health and entails ensuring reliable access to safe and clean water for human consumption, sanitation, and ecosystem functioning [2]. Several scientific studies have documented the existence of clinically significant antibiotic-resistant bacteria (ARBs) and ARGs in freshwater and wastewater [3–5]. However, wastewater (WW) systems have been reported to serve as a significant reservoir for diverse microbial communities, including multidrug-resistant (MDR) pathogens, contributing to the dissemination of AMR in the environment [6].

*P. aeruginosa* has been reported in sewage water and wastewater treatment plants (WWTP) sludge, consistent with levels found in hospital wastewater [7]. The increased prevalence of MDR-*P. aeruginosa*, particularly those resistant to the carbapenems, in sewage and WWTPs, emphasizes the urgent need for effective mitigation strategies [8–11]. Studies have demonstrated the persistence of MDR *P. aeruginosa* in treated effluents [5,12,13], downstream rivers [7,13], drinking water plumbing systems [14], and its dissemination through environmental pathways, thereby necessitating improved wastewater management. The presence of various virulence factors such as exopolysaccharide (EPS), flagella, and pili, intrinsic and acquired antibiotic resistance, and biofilm-forming potential confer this bacterium with a remarkable fitness and ability to adapt and thrive in various environments [15,16]. Conventional chemical treatments in WWTPs often fail to eradicate biofilm-associated bacteria and can contribute to the selection of resistant strains, perpetuating the cycle of AMR dissemination [17,18]. These limitations highlight the demand for sustainable alternatives capable of targeting biofilms and reducing the environmental burden of AMR [19].

Bacteriophages have resurfaced as eco-friendly biocontrol agents to address the issue of AMR and control microbial contamination in wastewater [20,21]. Several studies have explored the use of bacteriophages for reducing multidrug-resistant bacterial counts. However, these investigations mainly focused on the use of simulated WW with only the target bacteria [22], the treated WW [23], or the isolated bacterial strains from the WW to be treated independently with phages [24,25]. The unique biodiversity of phages within the Ganges River presents an opportunity to explore novel phages for environmental applications. Our recent study on the sediments of the river Ganges along its 1500 km has identified the repertoire of bacteriophages and their associated host-phage functions against putative human, plant, and putrefying pathogens [26]. However, there are limited studies with this riverine system wherein novel virulent phages were isolated from its waters against MDR strains of *Klebsiella pneumoniae* [24] and *P. aeruginosa* [25]. Our study aims to address these research gaps by isolating and characterizing ɸPRS-1 from the pristine (glacier-fed) stretch of the Ganges River. PRS-1 demonstrates efficacy in disrupting biofilms, downregulating *las* and *rhl* quorum sensing systems (QSS), and reducing the *Klebsiella pneumoniae* Carbapenemases (KPC) associated with MDR *P. aeruginosa*. By leveraging genomic and phenotypic analyses, this research seeks to advance our understanding of phage-based interventions in complex environmental matrices and contribute to sustainable WW management practices. The findings are anticipated to provide critical insights into phage therapy’s potential as a tool against AMR in WW, supporting environmental health, and achieving SDGs related to water security and public health.

## 2. Materials and Methods

### 2.1 Bacterial strain and antimicrobial susceptibility profiling

*Pseudomonas aeruginosa* PAO1 (ATCC 15692) (PAO1) was the bacterial host used. The genomic identity was ascertained with the MinION Mk1C Nanopore sequencer, Oxford Nanopore Technologies (ONT). The antimicrobial resistance profile of the isolate was determined via the VITEK 2 Compact (bioMérieux) using AST-N235 and AST-N281 cards. Antimicrobial susceptibility testing (AST) was performed according to Clinical and Laboratory Standards Institute (CLSI) guidelines.

### 2.2 Isolation, enrichment, and propagation of DPRS-1 targeting PAO1

Phage isolation against PAO1 (ATCC 15692) was carried out using sediment samples from Maneri (30°45’14“ N 78°33’34“ E), a pristine stretch of the Ganges River. The rationale was to explore and harness phages from this glacier-fed riverine environment. A distinct method was employed to isolate bacteriophages against PAO1. A recent study by Xuan et al. 2022, suggested that the QS machinery in PAO1 promotes phage infection wherein the bacterial lipopolysaccharide receptors required for the binding of the phages are upregulated. This phenomenon of *las*QS-mediated phage susceptibility, depending on the host cell density, increases the phage adsorption and the consequent infection of PAO1. The knowledge derived from the above study was employed to isolate virulent phages targeting PAO1 [27].

Sediment samples (n=3, 50 g each) were vortexed for 30 mins at room temperature. After settling, the sediment-laden water (50 mL) was equilibrated with 50 mL of Sodium-Magnesium (SM) buffer comprising NaCl (5.8g/L), MgSO_4_.7H_2_O (2g/L), Tris-HCl (pH 7.4) (50 mL), and distilled water (950 mL) for 4-5 hours at 37[and 120 rpm. The mixture was then centrifuged at 6000×g for 10 minutes, and the supernatant (∼50 mL) was filtered through 0.45μm and 0.22μm PES syringe filters. Ten milliliters of filtrate were used for phage enrichment against PAO1. A total of 5 mL of PAO1 culture in the log-stationary phase, grown in soybean-casein digest broth (SCDB), was mixed with 10 mL of filtrate and incubated at 37°C for 24 hours at 180 rpm. Lytic phages were confirmed by clear zones (plaques) in a qualitative spot assay. Phage enumeration was performed using the soft agar overlay method [28]. A single plaque was reconstituted in Sodium-Magnesium (SM) buffer, and its phage titer was determined. A singular, well-isolated plaque was serially diluted and infected with 100 µL of the log-stationary phase culture of PAO1 (6.5×10^8^CFU/mL). The host-phage mixture was plated using the double-layer agar plating method and incubated at 37 °C for 16-18 hours. Each set was titrated in triplicate across three independent iterations to ascertain the phage titer accurately. ɸPRS-1 was purified and concentrated using the Amicon® Ultra-15 centrifugal filter with a 100 kDa membrane. The multiplicity of infection (MOI) and Poisson distribution for predicting the probable number of infected cells in a population (P_0_) were calculated using the following formula [29]

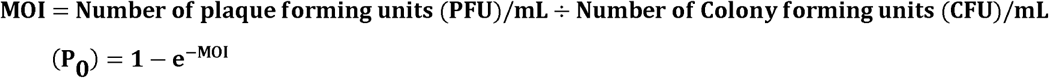

### 2.3 Characterization of PAO1 phage (**D**) PRS-1

#### 2.3.1 Morphological features

The morphology of ɸPRS-1 using 4 µL of high titer phage stock was visualized in Jeol JEM-F2100 high resolution-transmission electron microscope (HR-TEM) at 200 kV and imaged with a Xarosa emsis camera coupled to the microscope. To obtain a detailed (3D) view of the phage morphology, HR-TEM was used in STEM (Scanning Transmission Electron Microscopy) mode, enabling scanning images at nanometer (nm) resolution. Morphological classification of phage was done following the guidelines recommended by the International Committee on Taxonomy of Viruses (ICTV) [30].

#### 2.3.2 Phage adsorption rate and one-step-growth curve

The host cell culture at the exponential phase was mixed with the fresh phage lysate at an MOI of 0.1 and incubated at 37°C, 100 for 30 mins. After 5 minutes, a 1 mL sample was drawn and centrifuged to pellet the adsorbed fraction of the phage each time. At the same time, the supernatant was filtered using a 0.2 μm PES syringe filter to obtain the fraction of unadsorbed phages. The fractions were plated using the agar overlay method to get total phage (N _total_) and free phage (N _free_) densities, respectively. The adsorption rate (α) for each time point was calculated as:

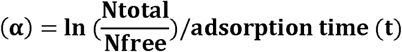

The determination of the one-step growth curve assay for ɸPRS-1 was carried out as described by Jagdale et al., 2019 with certain modifications [31]. Briefly, the fresh phage lysate of PRS-1 at an MOI of 0.1 was added to the PAO1 cells (10^8^ cells/mL) and allowed to adsorb for 15 mins at 37°C, 100 rpm. The adsorbed phages were centrifuged at 10,000 ×g for 5 min, and subsequently, the bacteria-phage complex (pellet) was resuspended in 2[mL of SCDB and incubated at 37°C, 100 rpm for 100 mins. A 100 µL sample was collected every 10[minutes, followed by plaque assay to determine the PFU/mL at each time point. The assay was carried out in triplicates.

#### 2.3.3 Evaluation of pH and temperature stability

The stability of ɸPRS-1 under different acidic and basic conditions was evaluated as described by (Oliveira et al., 2020) [32]. Briefly, 900 μL SM buffer with varied pH ranges adjusted to 3, 5, 7, 9, and 11 were inoculated with 100μL of the phage suspension (10^10^ PFU/mL), respectively, followed by 4 hours incubation at 37°C, 100 rpm in a dry bath, and the subsequent plaque assay. Similarly, for thermal stability evaluation, suspensions of 100 μL of the phage (10^10^ PFU/mL) were introduced in 900 μL of SM buffer and incubated at a diverse set of temperatures (4, 15, 25, 37, 45, 55, and 65°C) for 4 hours at 100 rpm followed by measuring the phage titer with plaque assay.

#### 2.3.4 Evaluation of the Lytic Spectrum of **D**PRS-1

The host range of ɸPRS-1 was evaluated using bacterial host cultures of *P. aeruginosa* (ATCC 15692), *P. aeruginosa* (ATCC 9027), *Escherichia coli* O157:H7 (ATCC 43888)*, Shigella boydii* (ATCC 9207, 8700)*, Salmonella enterica* (ATCC 12011, 13314)*, Staphylococcus aureus* (ATCC 6538)*, Listeria monocytogenes* (ATCC 19112)*, Enterococcus faecalis* (ATCC 19443) *Yersinia enterocolitica* (ATCC 27729). Details of the susceptibility profile of each host strain with ɸPRS-1 have been tabulated **(Table S1)**.

Besides this, the lytic spectra of ɸPRS-1 were evaluated on MDR *P. aeruginosa* isolates obtained from the sewage treatment plants (STP). For each of the *P. aeruginosa* isolates, the multiple antibiotic resistance index (MARI) was determined using the protocol followed by Cristina et al., 2021 [33]. After the qualitative spot assay, the virulence of phage was determined on MDR *P. aeruginosa* isolates through the efficiency of plating (EOP). In this, the phage titer at terminal dilution on the test strain was divided by the titer of the same phage on its host (isolation) strain. The virulence of фPRS-1 against the test strain was classified as highly virulent (EOP 0.1-1.00), moderately virulent (EOP 0.001-0.099), avirulent but active (EOP < 0.001), or avirulent (no plaques detected) [34], Details of *P. aeruginosa* specific host range, their MAR index, EOP, and virulence pattern have been tabulated in **Table 1**.

**Table 1:**
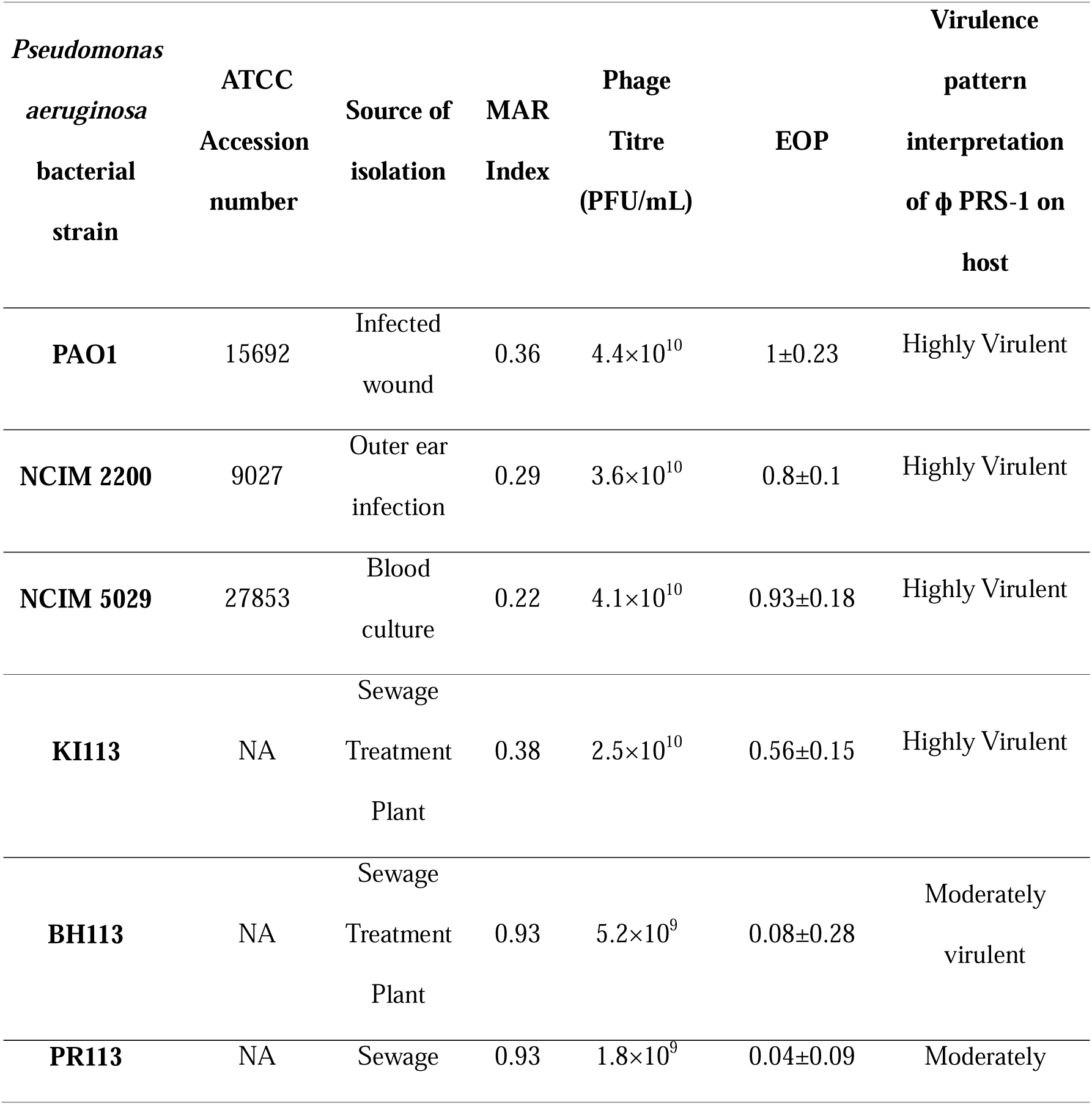
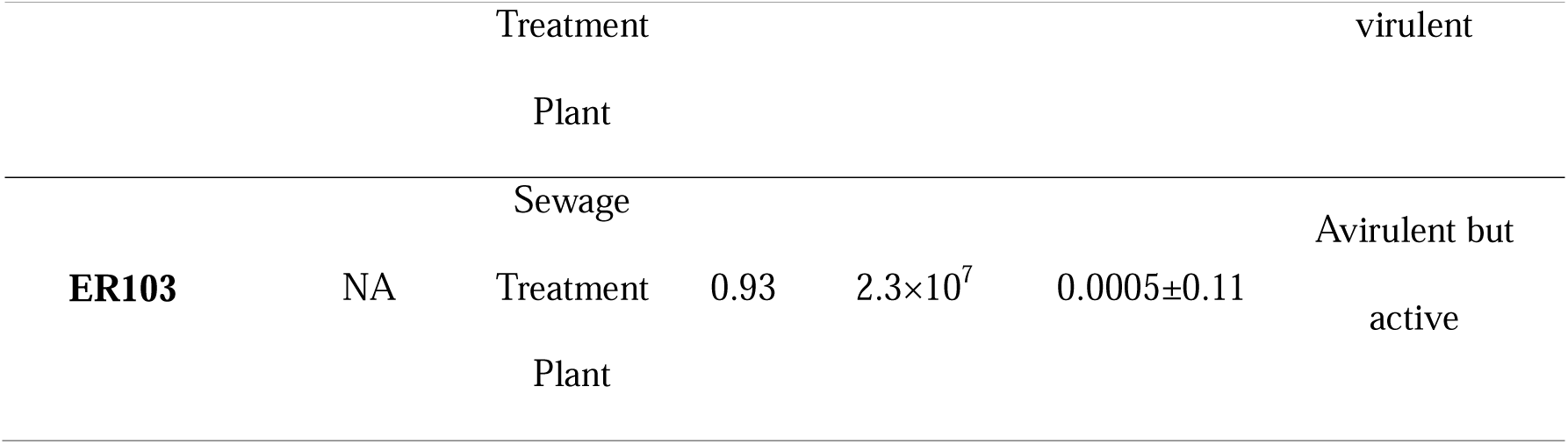
MAR index of *P. aeruginosa* strains and evaluation of the efficiency of plating (EOP) of phage PRS-1.

#### 2.3.5 DNA extraction, Genome Sequencing, and Analysis of **D**PRS-1

A high-titer phage stock was used to isolate DNA from ɸPRS-1. PureLink Viral RNA/DNA Mini Kit (Thermo Fisher Scientific) was used to extract the DNA, following the manufacturer’s protocol. The concentration of the phage DNA was measured on a Qubit 4 Fluorometer (Invitrogen). Whole genome library preparation was carried out using the Ligation Sequencing Kit (SQK-LSK114) and Native Barcoding Kit (SQK-NBD 114.24) with certain modifications **(Text S1)**. The library was then loaded onto a flow cell R10.4.1 (Oxford Nanopore Technologies) and sequenced for approximately 56 hours using the MinION Mk1C platform.

Sequencing reads were processed for quality control and adapter trimming using FastQC and Porechop [35]. Genome assembly was performed with Flye v 2.9.1 [36], and the assembled sequence was validated using NCBI nucleotide BLAST (BLASTN). Annotation of putative proteins was conducted with the RAST annotation server [37], and the resulting GenBank file was analyzed and visualized using Proksee [38]. The protein-coding sequences in the phage genome were identified with an open reading frame (ORF) finder from NCBI (https://www.ncbi.nlm.nih.gov/orffinder/).

Genome-wide similarities between ɸPRS-1 and reference viral genomes were analyzed using VipTree [39]. The average nucleotide identity (ANI) of the whole genome between ɸPRS-1 and closely related phages was computed with OAT v0.93.1[40]. The intergenomic similarities at nucleotide levels of ɸPRS-1 and other related viruses were determined with the VIRDIC tool [41]. Additionally, an integrated analysis of the phage’s identification, taxonomic classification, predictive lifestyle, and host was performed using PhaBOX [42]. The CARD database function in Proksee was used to analyze the presence of potential ARGs in the genome of ɸPRS-1.

### 2.4 Time-Dependent Evaluation of Antibacterial Efficacy of **D**PRS-1

The antibacterial efficacy of ɸPRS-1 was assessed against a thick coating of PAO1 cells (1×10^8^ CFU/mL) in square Petri plates (120×120 mm, Hi-Media). In this experiment, inhibition of bacterial growth by ɸPRS-1 was assessed as a function of time. A total of 10 mL of host culture was grown at the late log phase, followed by the addition of 0.0000001 MOI of phage. The CFU of PAO1 was determined at 0, 12, 24, and 48 hr in triplicates, and the CFU counts were determined at respective time points. Additionally, metabolic cell viability was assessed using resazurin, with relative fluorescence units measured at 550/590 nm.

### 2.5 Time-dependent Visualization and Quantification Antibiofilm Potentials of DPRS-1

The *in-vitro* efficacy of ɸPRS-1 against PAO1 (ATCC 15692) was evaluated for biofilm inhibition and disruption using the Film Tracer™ LIVE/DEAD® Biofilm Viability kit (Invitrogen).

#### Set I: Biofilm Inhibition

Herein, the overnight grown host culture cells were diluted in SCDB (1×10^8^ CFU/mL) and inoculated in a tissue culture dish, followed by the addition of ɸPRS-1 at 0.1 MOI. CFU was determined at 6, 12, and 24 hours, with live-dead cell imaging at 4, 8, 12, and 24 hours.

#### Set II: Biofilm Disruption

PAO1 cells were diluted in SCDB (1×10^8^ CFU/mL) and allowed to form a biofilm for 24 hours. After washing planktonic cells, ɸPRS-1 was added at 0.1 MOI. CFU was determined at 6, 12, and 24 hours with live-dead cell imaging at these respective time points. Untreated PAO1 cells served as controls in both sets.

For each time point, three replicates were used for confocal imaging, two for Field Emission Scanning Electron Microscopy (FE-SEM) sample preparation, and three for CFU determination. Cells were washed with 1X-PBS (Hi-Media) and stained with SYTO® 9 and propidium iodide (PI) from the Film Tracer™ LIVE/DEAD® Biofilm Viability kit, following the manufacturer’s protocol. Stained dishes were visualized using an inverted confocal laser scanning microscope (CLSM) (Leica Stellaris 5, DMi8) with a 20× objective, using 488 nm (SYTO^®^ 9) and 588 nm (PI) excitation wavelengths for live and dead bacteria, respectively. Phage PRS-1 treated and non-treated sets after 24 hours were also analyzed using (FE-SEM, Nova Nano SEM 450) to observe biofilm morphology changes.

### 2.6 Visualization and Quantification of Biofilm Disruption by **D**PRS-1 in single and Mixed Cultures of MDR-*P. aeruginosa* Isolates

A total of four MDR cultures of *P. aeruginosa,* designated as KI, BH, ER, and PR, were isolated from different STPs across Pune City in India. The antibiotics used during primary screening include Imipenem (10μg), Minocycline (30μg), Colistin (10μg), Levofloxacin (5μg), Ciprofloxacin (5μg), Aztreonam (30μg), Tigecycline (15μg), Cefoperazone/sulbactam (75/30μg), Ceftazidime (30μg), Cefixime (30μg), Co-Trimoxazole (25μg), and Piperacillin/Tazobactam (100/10). Details of their isolation and AST profiling have been comprehended in Supporting Information **(Text S2)**. Three of these MDR isolates (BH, KI, and PR), together with PAO1, were used for single and mixed culture biofilm experiments.

The overnight grown cultures of each MDR-*P. aeruginosa-*KI, *P. aeruginosa-*BH, *P. aeruginosa-*PR, *P. aeruginosa-*PAO1, and a mixture of all four strains (BH+KI+PR+PAO1) were diluted in SCDB (1×10^8^), inoculated in a tissue culture dish and allowed to form biofilm for 24-48 hours. The planktonic cells were washed, followed by the addition of ɸPRS-1 at 0.1 MOI. CFU was determined after 24 hours. Besides, the replicates were also subjected to live dead cell imaging at 6, 12, and 24 hrs.) as mentioned above in section 2.5. The untreated cells served as culture control in both sets.

### 2.7 Effect of **D**PRS-1 on Quorum Sensing and Carbapenemase Expression of MDR-*P. aeruginosa* isolates

The expression of quorum sensing systems (QSS) *las* and *rhl* was assessed using real-time reverse transcriptase polymerase chain reaction (qRT-PCR). The control set comprised the MDR-*P. aeruginosa* cultures (BH, KI, PR, PAO1) grown in SCDB broth till their late-log phase. In the treated set, 100 µL of ɸPRS-1 was added at 0.1 MOI to the cultures past their late-log phase and then incubated for 4 hours. RNA extraction was carried out using the RNeasy Powersoil kit (Qiagen). A normalized concentration of 100 ng/RNA μL was utilized for the preparation of cDNA with SuperScript IV VILO Master Mix (Invitrogen). Primers targeting QSS *lasI-R*, *lasR*, *rhlI*, *rhlR* [43], and *P. aeruginosa-*specific variable regions of the 16 S rRNA gene as the housekeeping gene [44] were used. The effect of phage on the key QSS was quantified in real-time using 1 µL of the cDNA with QuantiTect SYBR Green PCR Kit (Qiagen) following the manufacturer’s instructions. Additionally, the fate of specific carbapenemase genes (KPC, NDM, VIM, IMP, OXA[23, OXA[48, OXA[51, OXA[58), and their ARG concentration in the control and phage-treated group was quantified using Hi[PCR® Carbapenemase Gene Quantification Probe PCR Kit (Hi-Media) [45].

### 2.8 Time-dependent evaluation of *P. aeruginosa* biocontrol by **D** PRS-1 in untreated raw sewage sludge

A total of 1 L (n=3) of the raw sewage samples with sludge were collected from an STP, India (18° 32.8246’[N, 73° 48.8338’[E) in a sterile bottle by grab-sampling method. The experimental sets were divided into test and control groups. For the test group (100 mL aliquot of the raw sewage was challenged with 1 mL of high titer phage stock of [PRS-1(10^10^ PFU/mL), while the control group was devoid of any treatment. The experiment was carried out in triplicates, and the flasks from each group were incubated at 37°C for 24 hours. A time-kill assay was conducted to assess the potential of [PRS-1. The results were recorded as CFU/mL plated on cetrimide agar, and the reduction in the bacterial counts was calculated as log_10_ reduction and percent reduction, as described previously by Bashir et al. 2022 [46].

### 2.9 *In-vitro* Cytotoxicity Evaluation of DPRS-1

The *in-vitro* cytotoxicity of ɸPRS-1 preparations was assessed on L929, Vero mammalian cell lines, and Primary Human Umbilical Vein Endothelial Cells (HUVEC) using the (3-[4,5-dimethylthiazol-2-yl]-2,5 diphenyl tetrazolium bromide) (MTT) method described by Khan et al. 2024 [47]. A purified stock of ɸPRS-1 preparations, i.e., the enriched filtrate (∼2×10^8^ PFU/mL), high titer stock (3.8×10^10^PFU/mL), PEG precipitated stock (2.9×10^10^PFU/mL), endotoxin removed (ETR) preparation (2.5×10^9^PFU/mL), vehicle control (only SM buffer), and growth control (non/untreated cells) were evaluated for cytotoxicity on L929, Vero mammalian cell lines, and Primary Human Umbilical Vein Endothelial Cells (HUVEC).

Initially, the cells were propagated in T-flasks (Tarsons) using Dulbecco’s Modified Eagle Medium (DMEM) medium supplemented with 10% Fetal bovine serum (FBS) (Gibco). The confluency of the cells was ascertained through microscopic observation (Olympus CKX53 inverted microscope) of the cells. After achieving more than 80% confluency, the spent media was decanted from the cell culture flask. Subsequently, 200 μL of 1X Trypsin-EDTA solution (Hi Media) and an equal volume of DMEM were added to the flask and incubated for 2-3 mins in a CO_2_ incubator (Thermo Fischer Scientific). The contents were then transferred into a sterile 1.5 mL microfuge tube (Tarsons) and centrifuged at 6000 rpm for 2 mins. The supernatant was decanted, and the cells were resuspended in 1 mL of fresh DMEM medium.

From the resulting suspension, the cells were taken as 10 μL cells + 10 μL trypan blue + 80 μL PBS in separate sterile 200 μL microfuge tubes, and the number of cells was numbered using a Neubauer counting chamber. The seeding of the 96-well microtiter plate was done using 10^4^ cells in each well, followed by incubation at 37°C for 24 hours in a 5% CO_2_ incubator. Subsequently, phage lysates were added to the wells (>80% confluency) and incubated for 24-48 hours to assess the cytotoxicity. After 24 hours, the plate was visualized microscopically for morphological changes in the cells and incubated further for up to 48 hours. A total of 30 μL of MTT from 1 mg/mL stock was added to the wells. The plate was incubated in the dark for 4 hours at 37°C. Subsequently, the contents of the wells were decanted, and the resulting formazan crystals were solubilized with 100 μL DMSO. The resulting suspension was incubated for 5 mins, and the absorbance was recorded at 570 nm using a microplate reader (Biotek synergy H1).

The percent cell viability was calculated using the standard formula:

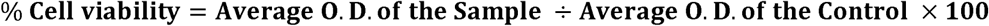

### 2.10 Statistical Analysis

The statistical significance of the assays was computed in GraphPad Prism v9.5.1 using repeated measures (RM) one-way Analysis of variance (ANOVA). Additionally, multiple comparisons with a false discovery rate (FDR correction) and a p-value of 0.05 were used to compare the mean between the groups as a function of time.

### 2.11 Data Availability

The genome sequence data of ɸPRS-1 has been submitted to GenBank (BankIt tool) under accession number PP337210. The SRA Biosample and Bioproject accession numbers for ɸPRS-1 are SAMN41895092 and PRJNA1125352. The annotated data of ɸPRS-1 has been deposited in the Zenodo repository with the DOI <u>10.5281/zenodo.12097649</u>.

## 3. Results and Discussion

### 3.1 Isolation, identification, and growth characteristics of DPRS-1

A recent study by Xuan et al. 2022, suggested that *las*QS-mediated phage susceptibility, depending on the host cell density in the late-log phase, increases the phage adsorption and the consequent infection of PAO1. The knowledge derived from the above study was employed to isolate virulent phages targeting PAO1 (ATCC 15692) [27]. This enrichment method resulted in the isolation of 12 phages targeting PAO1. Amongst all the isolated phages, ɸPRS-1 showed significant inhibitory activity against PAO1 for 48 hours **(Fig 1A)**. Consequently, ɸPRS-1 was chosen further for detailed characterization in this study. The host range of ɸPRS-1 has been tabulated **(Table S1)**.

**Figure 1:**
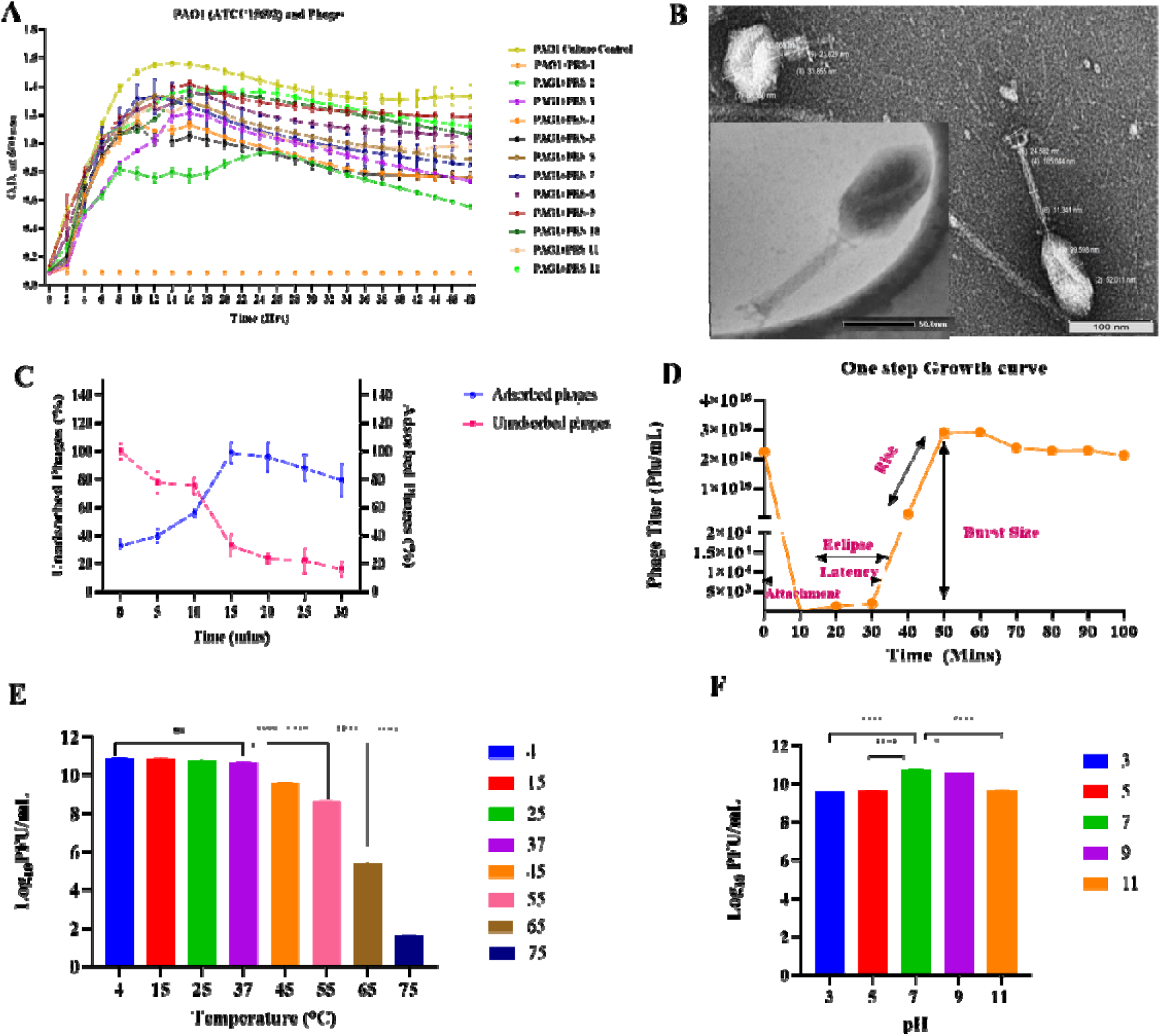
Morphology and growth parameters of phagePRS-1. **(A)** Screening of potent phages against PAO1. **(B)** HR-TEM micrographs of phagePRS-1 in its extended and contracted state at 80000× and scalebar of 100 nm. The insets feature- top right: plaque morphology, and bottom left: a STEM image of phagePRS-1 in its extended state. **(C)** A graphical illustration of the variation in the percentage of adsorbed and unadsorbed phage particles of phagePRS-1 as a function of time. **(D)** One-step growth curve of phagePRS-1 with the phases of its lifecycle marked (pink). **(E)** A graphical illustration depicting the influence of temperature on the stability of phagePRS-1 **(F)** A graphical illustration demonstrating the impact of various pH ranges on the stability of phagePRS-1. The data depicted in graph panels C through F are the average values of experimental replicates and their standard deviations (SD). The symbols (*, **, ***, ****) are indications of statistical significance determined using repeated measures of one-way ANOVA (RM-ANOVA). No significant difference between the compared test groups is designated with the abbreviation ‘ns.’

The AMR profile of PAO1 through VITEK-2 indicated its resistance toward more than three antimicrobial classes, namely the members of second and third-generation cephalosporins (2GC, 3GC), beta-lactams, quinolones, second and third-generation tetracyclines, diaminopyrimidine, phosphonic and nitrofuran class of antibiotics. The detailed AST profiles for PAO1 with the MIC values have been summarised **(Supporting information*_*PAO1*-*AST281, PAO1-AST235)**. Additionally, the MARI of PAO1 was recorded as 0.36, indicating that the PAO1 strain has been exposed to several antibiotics or disinfectants. Therefore, the use of a novel bacteriophage to tackle the MDR *P. aeruginosa* is an effective alternative strategy amidst the AMR crisis.

The genome annotation of PAO1 highlights several MDR mechanisms, including MexEF-OprN efflux pumps and metallo-beta-lactamases (MBLs), such as carbapenemases, emphasizing the need for phages devoid of ARGs to manage MDR-*P. aeruginosa* effectively. The details of genome annotation, with particular emphasis on the MDR profile of PAO1, have been summarised in supporting information **(Table S2)**.

The MARI analysis of PAO1 alongside other MDR *P. aeruginosa* isolates from sewage treatment plants (STPs) revealed MARI values >0.2 for all isolates, indicating substantial exposure to antibiotics or disinfectants. Furthermore, the AMR profiling showed that these seven isolates are resistant to more than three classes of antibiotics, indicating their MDR nature. This underscores the need for novel phages like ɸPRS-1 against MDR *P. aeruginosa* in wastewater. Details of the *P. aeruginosa-*specific host range, their MAR index, EOP, and virulence pattern have been tabulated in **Table 1**.

The ɸPRS-1 comprised large clear plaques surrounded by a halo, which could be attributed to the presence of polysaccharide depolymerase, either virion-associated (the structural proteins or tail fibers) of ɸPRS-1 or of the soluble type, which diffuses in the surrounding medium [48]. Morphologically, ɸPRS-1 exhibits myovirus-like features. The structure of ɸPRS-1 comprised of a prolate head (length, 99 ± 2 nm; width, 52[±[3 nm) showing an icosahedral symmetry, a collar with whiskers, a long contractile tail (length, 105[±[5 nm; width, 11[±[2 nm), a small baseplate (24 ±[2 nm) with short spikes, and around six terminal (tail) fibers **(Fig 1B)**. Based on these morphological characteristics, ɸPRS-1 was classified within the *Caudoviricetes* class as per the International Committee on Taxonomy of Viruses (ICTV) [30].

The titer of ɸPRS-1 was determined as 4.1×10^10^ PFU/mL, having an MOI of 6.30, and effective in achieving a bacterial clearance of 99.81%, with a probability of infected cells being P_0_=0.9981. An MOI of five or higher suggests a simultaneous infection of the host cell by more than one phage particle, indicating the virulent ability of the phage to establish a productive infection within the susceptible host [49].

The adsorption studies with ɸPRS-1 indicated >90% by 15 mins with a simultaneous decrease in the percentage of free phages in the unadsorbed fraction **(Fig 1C)**. Details of the adsorption rate have been compiled **(Table S3)**. Further, the replication cycle of ɸPRS-1 showed a latent period of ∼25 mins and a burst size of 37 (±5) phage particles per infected cell of PAO1 (host) **(Fig 1D)**. Interestingly, optimal temperatures for phage proliferation were observed from 4[to 37[with no significant change in the phage titer. This phenomenon may be attributed to the origin of the phage, which is from the upper stretch of the Ganges River at the Maneri location, where temperatures are relatively colder. However, a notable decrease in the phage titer was observed when the temperature increased from 37[to 75[(ANOVA, p<0.001). **(Fig 1E)**. Based on the phage titration, pH 7 was identified as the optimal pH for maintaining phage stability. On either side of the pH scale, it was observed that the phage titer was reduced **(Fig 1F)**. However, a phage titer of 10^9^ PFU/mL is considered a high titer [50]. This suggests that ɸPRS-1 continues to have an infectious nature for PAO1 even at a diverse pH range. The distribution of *P. aeruginosa* is ubiquitous, and it persists in environments with varied temperatures and pH [51–53]. The experimental evidence suggests that ɸPRS-1 can infect its target host across different temperatures and pH, making it an ideal candidate to tackle the MDR *P. aeruginosa* in different types of wastewater.

### 3.2 Genome analysis, annotation, and proteome of **D**PRS-1

The genome of ɸPRS-1 spans 65,628[bp with a 57.6% G+C content. Analysis of open reading frames (ORFs) identified 799 ORFs, primarily consisting of hypothetical proteins, undefined phage proteins, and proteins that are responsible for defining phage structure, DNA replication, repair, lytic cassette, and packaging of the virion. The specific positions and details of various proteins within the genome of ɸPRS-1 have been depicted in **Fig 2A**. From genome annotation using RAST and Prokka v1314.6 tool, endolysin was identified as the key enzyme responsible for the host lysis by ɸPRS-1.

**Figure 2:**
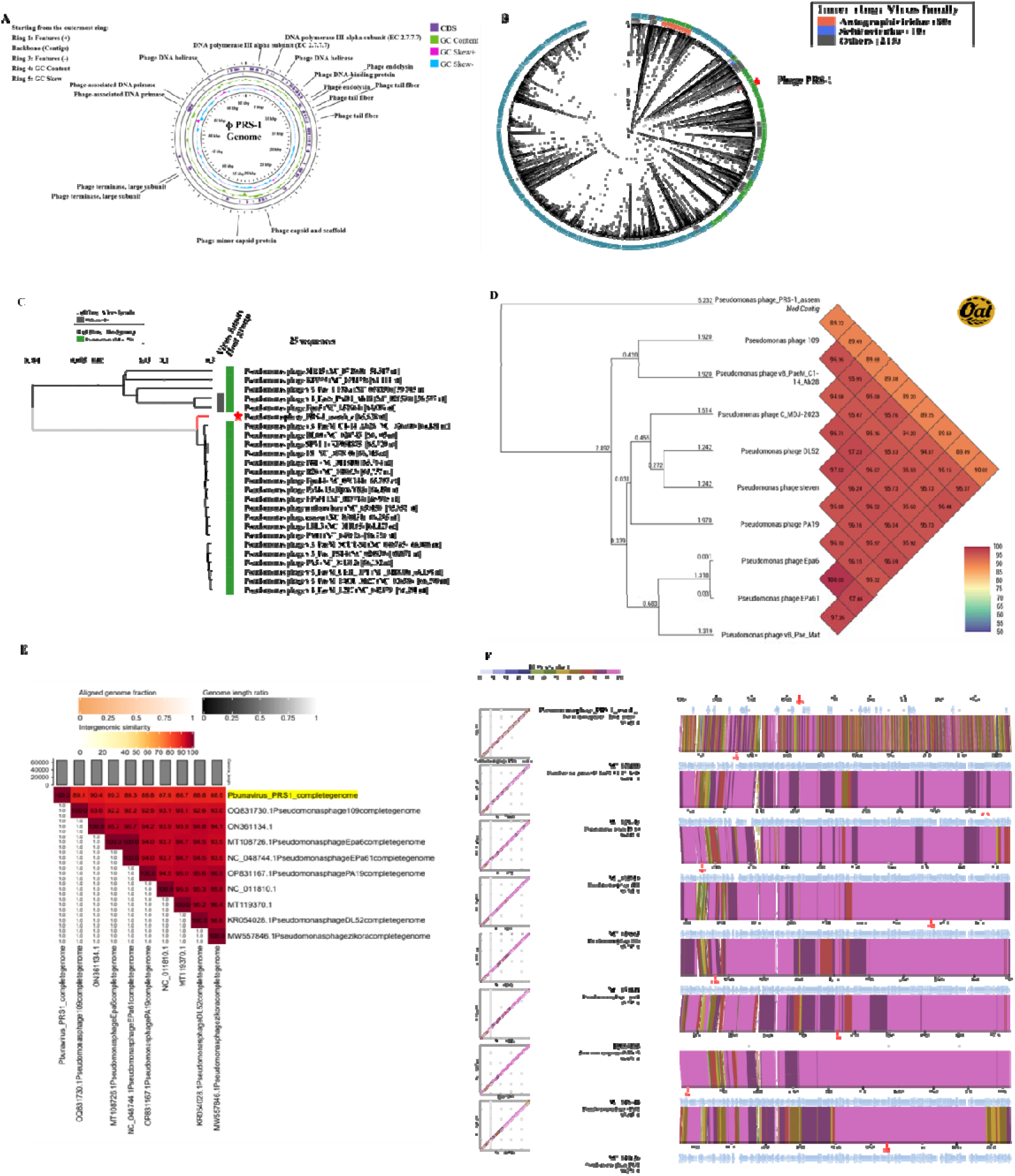
Genomic Features and Amino AcidBased Comparative Assessment of phage PRS-1. **(A)** A circular genome map of phagePRS-1 was generated using Proksee viewer. The genome contents are shown as ring diagrams. The outer rings in purple arrows represent the RAST-annotated protein-coding genes of the positive and negative strands of the DNA. The inner circle in green indicates the total GC content, while the pink and blue colors depict GC skew information for the positive strand and negative strands, respectively. **(B)** A circular phylogenetic tree, created from VipTree, with 1997 reference genomes of phages related to phagePRS-1. The genome distance matrix between phagePRS-1 and its relatives is computed using BIONJ, an algorithm based on the distance for phylogeny reconstruction. The midpoint of the tree indicates the root, with outer-colored rings representing host taxa and inner-colored rings representing viral families. **(C)** A rectangular phylogenetic tree of phagePRS-1, along with its 20 closely related and 4 distantly related phage genomes, constructed based on S_G_ scores. The red star denotes Pseudomonas phage phagePRS-1. **(D)**. UPGMA tree illustrating OrthoANI values between phagePRS-1 and closely related *Pbunaviruses* generated using OAT software. The color scale (lower right) represents the similarity percentage between the genomes. The numerical values on the branches indicate branch length as determined by the OAT software. **(E)**. The heat map shows alignment indicators (left half) and intergenomic similarity values (right half) generated through VIRDIC. **(F)** A synteny plot generated by TBLASTX using the ViPTree server illustrates whole genome sequence alignment phage PRS-1 with eight closely related phages. The light blue arrows represent the open reading frames (ORFs). The color of the bands between the genetic maps indicates the % identity from TBLASTX, as specified in the legend at the top of the figure.

Analysis of amino-acid sequences in the whole genome of ɸPRS-1 and closely related phages was performed using VipTree. A total of 1997 genomes related to ɸPRS-1 were used to construct the phylogenetic tree **(Fig 2B)**. The initial observation of the circular phylogenetic tree of ɸPRS-1 was consistent with the morphological features of myovirus. Furthermore, based on the genomic similarity (SG) scores from tblastx, a subset comprising twenty closely related phages and four distantly related phages was selected to construct a rectangular phylogenetic tree. **(Fig 2C)**. Further, from the orthoANI comparison of the genome of ɸPRS-1 and closely related *Pbunavirus* members, it was observed that ɸPRS-1 is 90.02% related with Pseudomonas phage vB_Pae_Mat and 89.72% with Pseudomonas phage 109, indicating the close relatedness of its genome as evident in the unweighted pair group method with arithmetic mean (UPGMA) tree **(Fig 2D)**. As per ICTV, the isolated phages are classified within the same genus if the identities of their nucleotide sequences exceed 70% [54]. This indicates that ɸPRS-1 could be classified as a new member within the unclassified family of the *Caudovircetes* class and *Pbunavirus* genus.

Additionally, a whole genome comparison was made by aligning the genome of ɸPRS-1 and eight closely related phages based on the genome similarity (S_G_) scores **(Fig 2F)**. The findings revealed differences in the synteny of ɸPRS-1 compared to the related phages, indicating its separation as a distinct member within this group of phages.

Furthermore, PhaBOX Analysis of the phage lifecycle and CARD analysis of the phage genome revealed that the ɸPRS-1 is an absolute lytic phage (devoid of lysogeny-related genes) and shows no apparent presence of any antimicrobial resistance genes in its genome **(Supporting Information PhaBOX results)**.

The members of *Pbunavirus* are known to infect *Pseudomonas* species exclusively. (https://www.ncbi.nlm.nih.gov/Taxonomy/Browser/wwwtax.cgi). However, it is important to note that there are no reports in the NCBI database of the phages from the *Pbunavirus* genus isolated from the Ganges River and that which infects *P. aeruginosa*. Therefore, to the best of our knowledge, this study represents the first comprehensive report providing a detailed analysis of the complete genome, as well as nucleotide and amino acid-based phylogenetic understanding, of a *Pbunavirus* member isolated from an unexplored source of the Ganges River, having the capability to infect MDR *P. aeruginosa* strains (ATCC 15692, 9027).

### 3.3 Antibacterial potential of **D**PRS-1

The antibacterial efficacy of ɸPRS-1 was assessed at various dilutions, with the lowest effective MOI being 10^-7^ **(Fig 3A)**. The assessment of cell viability after phage treatment indicated a remarkable reduction of 8.14 log_10_ with a percent reduction of >99.99% in CFU/mL of PAO1 at 10^-8^ dilution **(Fig 3B)**. Additionally, time-dependent evaluation of the antibacterial effect of ɸPRS-1 at 12, 24, and 48 hours was evaluated with PAO1 cells. The results showed a remarkable 6.42 log_10,_ 7.35 log_10,_ and 8.67 log_10_ reductions at 12, 24, and 48 hours. Additionally, a lesser resazurin signal in the ɸPRS-1 treated sets is indicative of less metabolically active cells, thereby supporting the antibacterial efficacy of ɸPRS-1 (**Fig 3C, D)**. Several studies with MDR *P. aeruginosa* and phage(s) in diverse sets of environments have shown a 1.5-5 log_10_ reduction [55–57]. The commercially available *Pseudomonas* phage JG003 shows a 1.2 log_10_ reduction in PAO1 cell count [58]. However, to our knowledge, this is the first report showing >8 log_10_ reduction in the viable count of MDR pathogen PAO1 from 0.0000001 MOI of a single, non-engineered phage.

**Figure 3:**
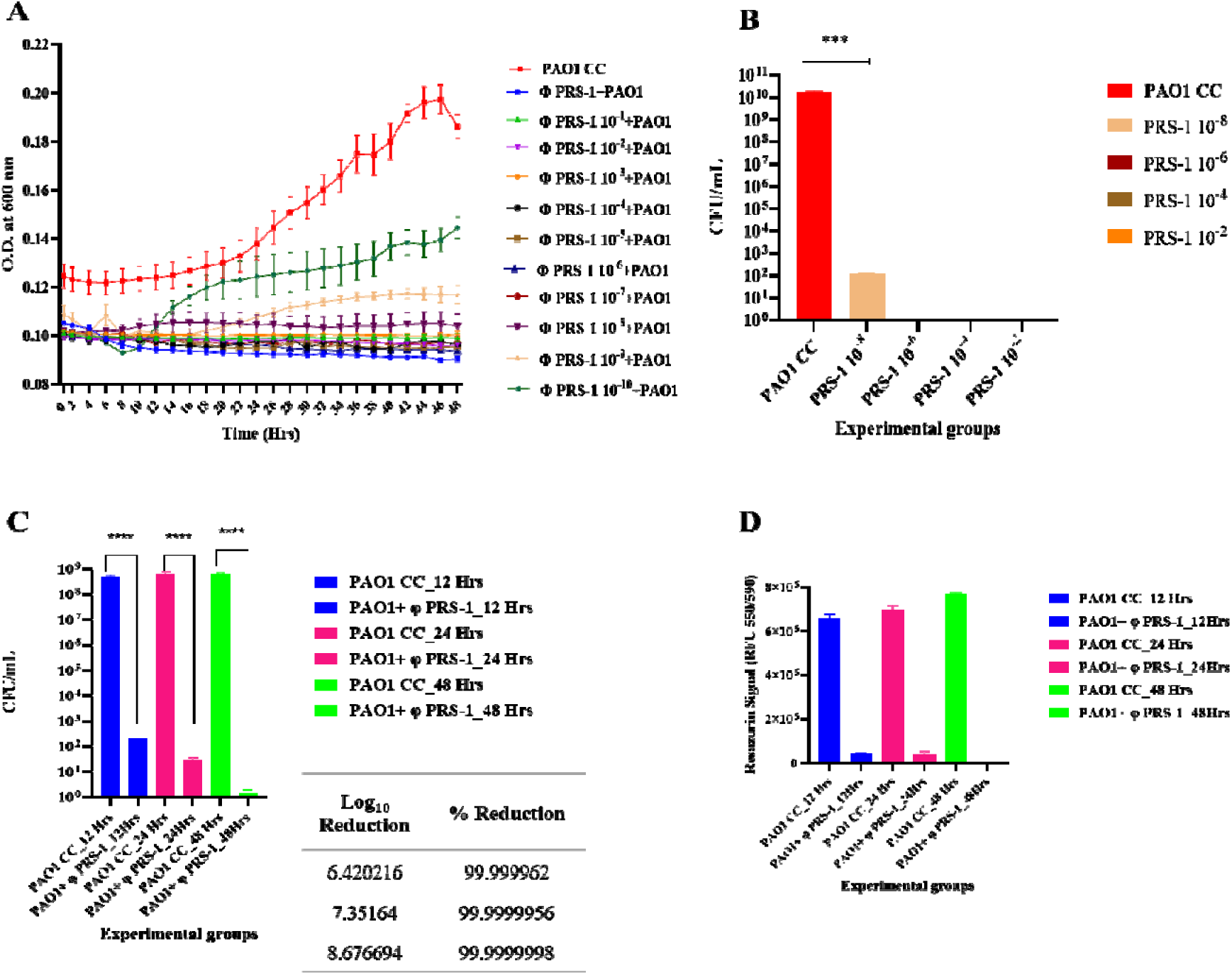
Antibacterial efficacy of phage PRS-1. **(A)** OD _600-_based evaluation of the inhibitory effect of various dilutions of phagePRS-1 against *P. aeruginosa* PAO1. **(B)** Quantitative estimation of cell viability (CFU/mL) of PAO1 cells after treatment with various dilutions of phagePRS-1**. (C)** Time-dependent evaluation of the antibacterial effect of phagePRS-1 on the cell viability of PAO1. **(D)** Time-dependent assessment of metabolic cell viability of PAO1 cells treated with phage PRS-1.

### 3.4 Time-dependent evaluation of *P. aeruginosa* PAO1 biofilm inhibition and disruption

The genome annotation of PAO1 revealed its biofilm composition to be primarily composed of a dimeric repeat of α-1,4 linked galactosamine and N-acetylgalactosamine, required to maintain the structural integrity of the biofilm **(Table S2)**. The CLSM analysis of ɸPRS-1 treated PAO1 set showed red fluorescence at 4, 8, 12, and 24 hours, indicating the dead biomass of PAO1 cells **(Fig 4 B-E)**, with a remarkable 7.8 log_10_ reduction at 4 hours reaching 9.83 log_10_ at 24 hours (ANOVA, p=<0.0001) **(Fig 4J)**, as compared to the control set **(Fig 4A)**. The FE-SEM analysis corroborated these findings, showing aggregates of dead cells with ruptured cell membranes, indicating cell lysis by ɸPRS-1 **(Fig 4H-I)**. The biofilm matrix of the control set was found to be intact **(Fig 4F)** with a typical coccobacillus morphology of *P. aeruginosa* cells **(Fig 4G)**. Several studies have implemented the use of bacteriophages in biocontrol of MDR *P. aeruginosa* biofilms on various fomites, medical implants, and tubings, *in-vivo* studies demonstrating the effect of phages in controlling *P. aeruginosa* biofilms in various organs with a reduction of 2.5-5 log_10_ within 24 hours [57,59,60]. However, this is the first report with phage signifying a remarkable reduction of 9.83 log_10_ of MDR *P. aeruginosa* in its biofilm form within 24 hours.

**Figure 4:**
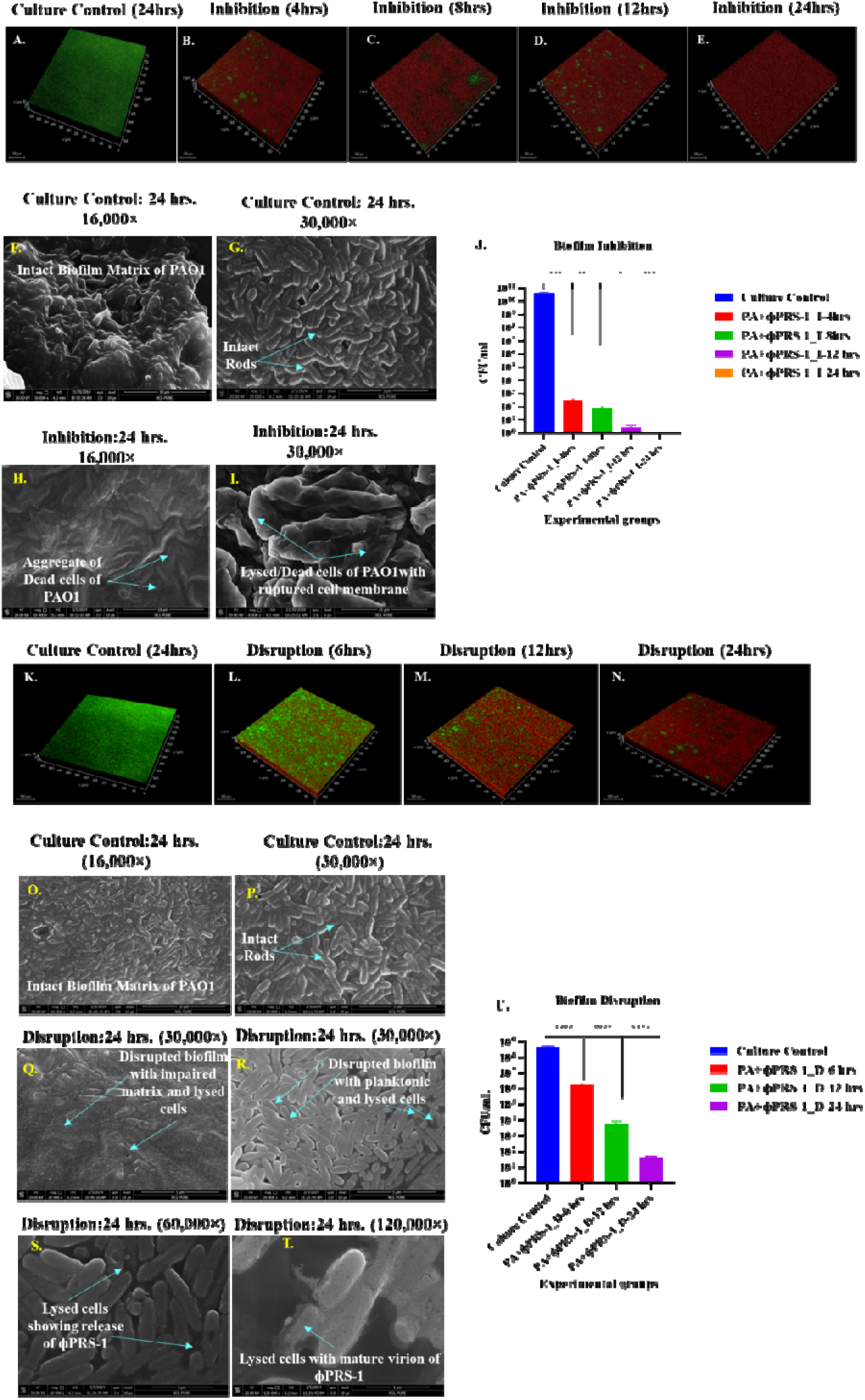
Effect of phage PRS-1 on biofilm formation and disruption by *P. aeruginosa* PAO1. Samples were observed under a Leica Stellaris 5, DMi8 microscope with a 20× objective. Dead *P. aeruginosa* PAO1 cells are indicated by red fluorescence, while viable cells are indicated by green fluorescence at various time intervals. **(A)** CLSM image of untreated *P. aeruginosa* PAO1 cells capable of forming biofilm after 24 hours. **(B)** CLSM images of the phagePRS-1 treated group were captured after 4 hours **(B)**, 8 hours **(C)**, 12 hours **(D)**, and 24 hours **(E)**. **(F-I)** FE-SEM images of biofilm. Culture control showing intact rods embedded within the EPS matrix of the biofilm 16,000×**(F)** 30,000× **(G)**. **(H-I)** Biofilm Inhibition by phage PRS-1 showing dead and lysed cells of PAO1. **(H)** Aggregate of dead cell biomass at 16,000×. **(I)** A magnified view of dead PAO1 cells with ruptured cell membranes. **(J)** Quantification of the viable cells (CFU/mL) in both control (non-treated) and phagePRS-1 treated sets. **(J)** CLSM image of culture control with preformed biofilm of PAO1 cells (lacking any treatment). **(K-N)** CLSM micrograph of phage PRS-1 treated preformed biofilm imaged after 6 hours **(L)**, 12 hours **(M)**, and 24 hours **(N)**. **(O-T)** FE-SEM images of control biofilm at different magnifications 16000× **(O)**, 30000× **(P)**. Disrupted matrix of the PAO1 biofilm 30000× **(Q)**. **(R)** Disrupted biofilm with lysed planktonic cells. **(S-T)** Lysed cells of PAO1 show the release of the progeny of phage PRS-1. 60000× **(S)**, and 120000× **(T)**. **(U)** Quantification of the viable cells in disrupted biofilm as CFU/mL of the control and phagePRS-1 treated sets.

Biofilms of various microbes exist in their mature state in the environments. Thus, to detach the biofilm, it is crucial to break down its EPS [55]. The results indicated a gradual decrease in the viable cells of the ɸPRS-1 treated group **(Fig 4 K-N, Fig 4U)**. Therefore, this reports the first evidence of a 7 log_10_ reduction within 24 hours in preformed biofilm using a single phage. The FESEM imaging showed a well-defined, rod-shaped morphology immersed in the EPS matrix of the control **(Fig 4 O, P).** In contrast, the ɸPRS-1 treated set showed an impaired EPS matrix with distorted morphology of the cells **(Fig 4 Q-T)**. The potential of ɸPRS-1 to inhibit and disrupt PAO1 biofilm could be attributed to the phage depolymerase enzyme, which cleaves the EPS matrix and disarms the bacterial virulence, explaining the clear plaque with a surrounding halo.

### 3.5 Visualization and Quantification of Biofilm Disruption by **D**PRS-1 in Single and Mixed Cultures of MDR *P. aeruginosa* Isolates

Four MDR *Pseudomonas aeruginosa* cultures, designated KI, BH, PR, and PAO1, were evaluated for biofilm disruption potentials of ɸPRS-1. Quantitative analysis (CFU/mL) of the biofilm cell viability results showed a marked decrease in biofilm CFU across all phage-treated samples compared to the untreated controls **(Fig 5A)**. For instance, BH+PRS-1 showed a reduction to 4.940714 log CFU/mL, with a viability of 99.9985%. The mixed culture (All 4+PRS-1) demonstrated a similar trend with a log reduction to 5.870824 and 99.9987% viability, further supporting the enhanced efficacy of ɸPRS-1 in disrupting complex biofilm structures **(Fig 5C)**. These observations align with the visual evidence of the phage’s impact through live-dead cell imaging. Here, green fluorescence denotes live cells, while red fluorescence indicates dead cells. Untreated biofilms across all isolates displayed dense green fluorescence, signifying high cell viability. In stark contrast, biofilms treated with LPRS-1 showed a substantial increase in red fluorescence, indicating extensive cell death and biofilm disruption. The mixed culture biofilm (All 4+PRS-1) displayed the most pronounced red fluorescence, aligning with the highest quantitative reduction observed **(Fig 8B)**.

**Figure 5:**
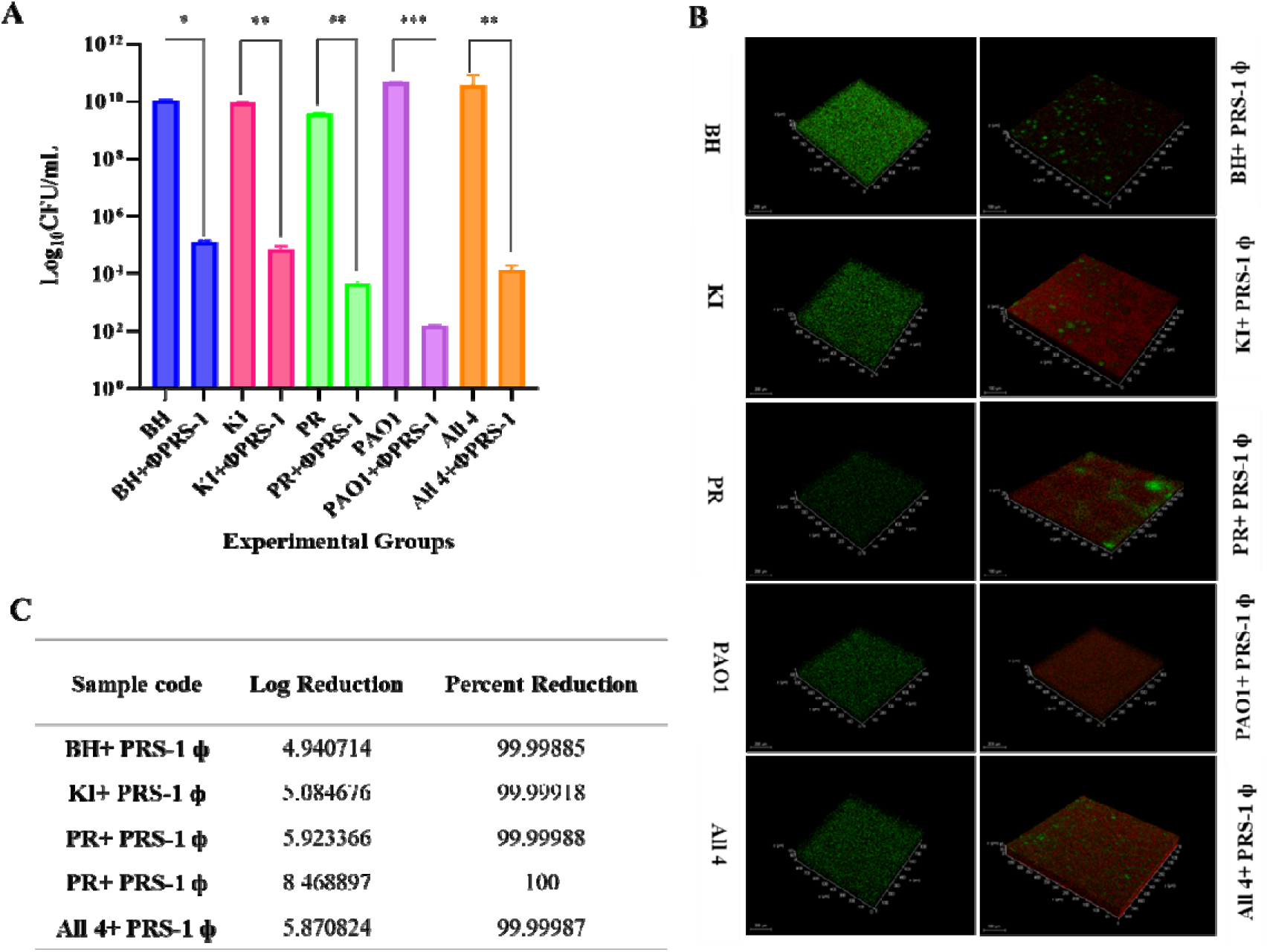
Quantitative Analysis and Live-Dead Imaging of Biofilm Disruption by Bacteriophage DPRS-1. **(A)** Bar graph showing the quantitative analysis of biofilm disruption in MDR *P. aeruginosa* isolates BH, KI, PR, and PAO1, as well as a mixed culture (All 4), after treatment with bacteriophage LPRS-1 at 0.1 MOI. **(B)** Live-dead cell imaging of biofilms using fluorescence microscopy. Images display the untreated and LPRS-1 treated biofilms of MDR *P. aeruginosa* isolates BH, KI, PR, PAO1, and the mixed culture (All 4) at 24 hours post-treatment. **(C)** Quantification of biofilm CFU and cell viability after treatment with ɸPRS-1.

### 3.6 Effect of **D**PRS-1 on the expression of *las* and *rhl* quorum sensing systems and carbapenemases

In this study, we demonstrated that ɸPRS-1 disrupts the regulatory networks by downregulating the QS genes (fold decrease): *lasI-R* (−2.6), *lasR* (−3.4), *rhlR* (−6.3), and *rhlI* (−3.1), in the *las* and *rhl* QSS, essential for *P. aeruginosa* virulence and biofilm formation **(Fig 6 A, B)**. Our results align with previous findings showing phages can reduce las and rhl gene expression, thereby decreasing virulence and biofilm formation [61,62]. While prior research has established phages’ potential to modulate QS systems, direct evidence of phage-induced downregulation of carbapenemases like KPC remains limited. Notably, ɸPRS-1 markedly reduced KPC carbapenemase gene copy numbers in MDR-*P. aeruginosa* isolates: from 6,841,800 to 80,187 in BH, from 3,267,790 to 47,455 in KI, from 5,482,877 to 8,133 in PR, from 1,013,330 to 8 in PAO1, and all 4 collectively from 590,579 to 1,392 **(Fig 6 C)**. KPC concentrations (ng), determined using the standard graph, were reduced in the phage-treated counterparts to 9.2, 8.4, 4.8, 0.2, and 1.6 ng, respectively, from control values of 16, 15, 14, 12, and 12 ng **(Fig 6D)**. These findings expand our understanding by providing the first documented evidence of phage-mediated reduction of carbapenemases in MDR *P. aeruginosa*, highlighting the potential of ɸPRS-1 in abatement carbapenemase resistance in this WHO priority pathogen. This marks the first experimental validation of a bacteriophage effectively tackling the MDR *P. aeruginosa* AMR through the targeted downregulation of both *las* and *rhl* QSS and reduction of carbapenemases.

**Figure 6:**
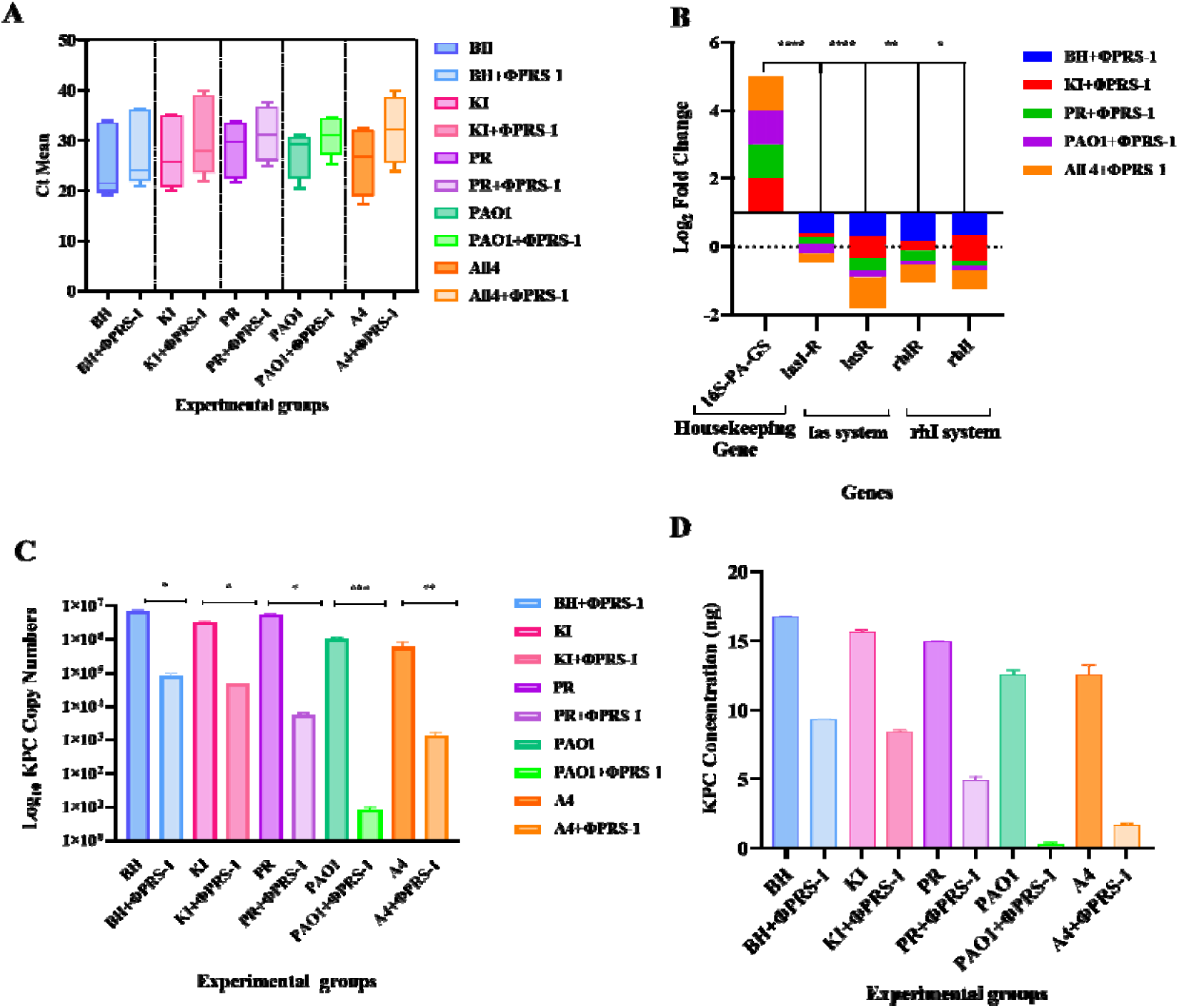
Effect of DPRS-1 on the expression of *las* and *rhl* quorum sensing systems and carbapenemases. **(A)** Graphical illustration of Ct means of the LPRS-1 treated and control groups for assessing the effect on the expression of *las* and *rhl* quorum systems. **(B)** Log_2_ fold change in the expression of *las* and *rhl* quorum systems sowing downregulation in LPRS-1 treated sets. **(C)** Reduction in the copy numbers of KPC carbapenemase gene expression in LPRS-1 -treated sets. **(D)** Quantification of the concentration of the KPC in the control and LPRS-1 -treated groups.

### 3.7 Time-dependent evaluation of reduction in *P. aeruginosa* counts from raw sewage sludge

Treatment of raw sewage with LPRS-1 significantly reduced viable *P. aeruginosa* counts with a 2.4 log_10_ reduction at 6 hours and a 4.2 log_10_ reduction at 24 hours (ANOVA, p<0.001) **(Fig 7 A, B)**. VITEK-MS-based characterization of the colonies that grew on the cetrimide agar plates of the ɸPRS-1 treated group revealed *Pseudomonas putida, P. luteola,* and *Serratia marcescens* to be among the survivors. However, at any given timepoint, strains of *P. aeruginosa* were not recovered from [PRS-1 treated group, indicating the efficacy of [PRS-1 in biocontrol of *P. aeruginosa-*specific strains from raw wastewater combined with the sludge.

**Figure 7:**
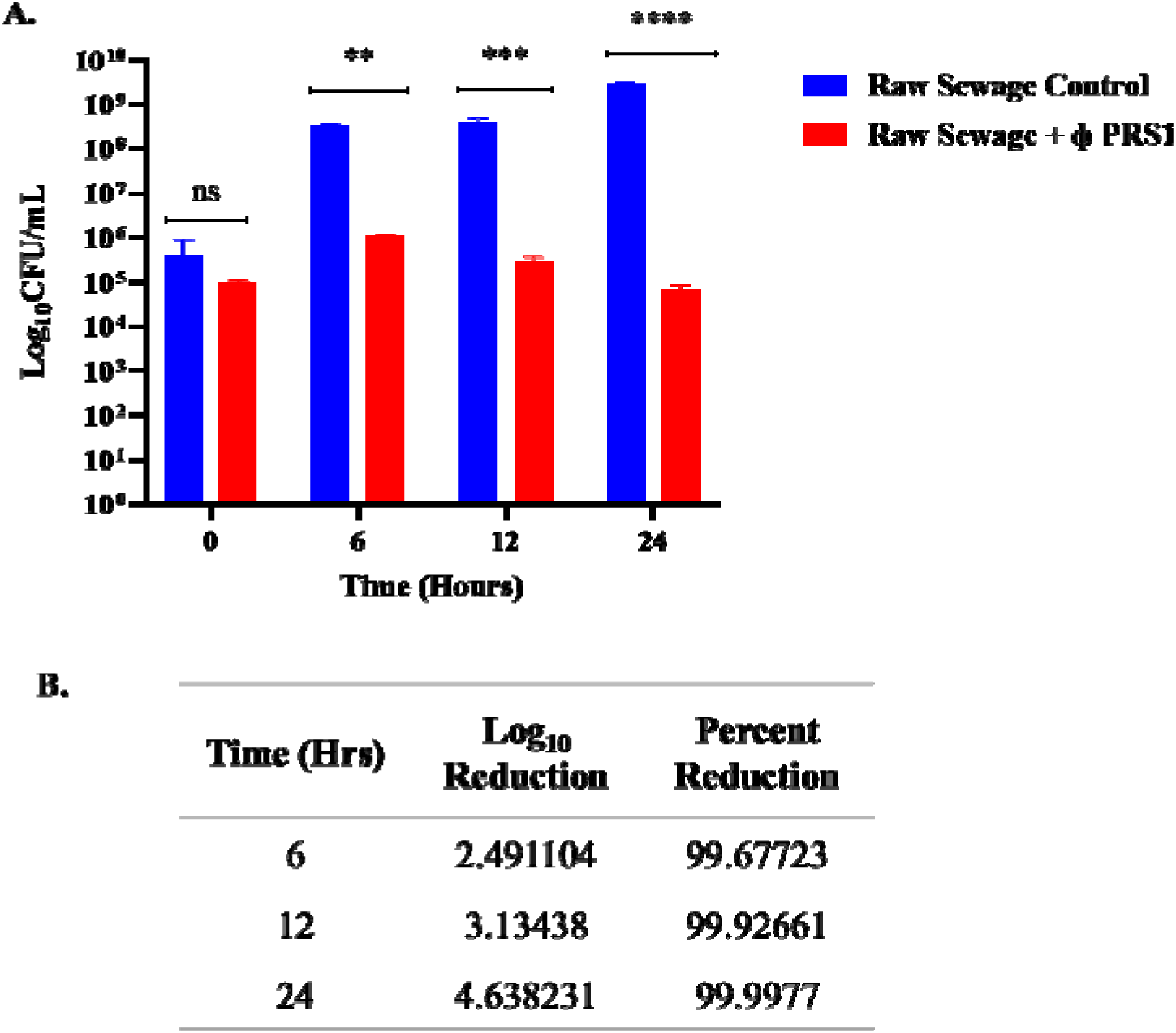
Evaluation of DPRS-1 in reduction of *P. aeruginosa* specific counts in raw sewage wastewater. **(A)** Graphical representation of quantification of the viable cells in raw sewage as CFU/mL of the control (non-treated) and L ERS-1 treated sets. **(B)** Log reduction and percentage reduction values indicate L ERS-1 efficacy in reducing bacterial load in raw sewage after 24 hours.

To date, a few studies have detailed the application of phages, in WW treatment to reduce pathogenic and MDR bacterial populations [23,63,64]. To our knowledge, this study is the first of its kind to detail an in-depth characterization of a novel ɸPRS-1, highlighting its antibacterial and antibiofilm capabilities, along with the lab-scale demonstration of its effectiveness in reducing MDR *P. aeruginosa* bacterial counts in raw sewage. Nevertheless, achieving a thorough understanding of the optimized phage dosage and its effects on physicochemical and microbiological parameters pre- and post-treatment with [PRS-1 is crucial for its practical application on-site.

### 3.8 *In-vitro* cytotoxicity evaluation of **D**PRS-1 on L929, Vero, and HUVECcell lines

As LPRS-1 is a novel phage, the knowledge of its cytotoxic effects is of crucial importance. The results showed no observable difference in the cellular morphology of the control cells versus those treated with LPRS-1 preparations when imaged after 48 hours of incubation in L929, Vero, and HUVECcell lines. **(Fig 8A-C)**. MTT assays indicated 95-99% cell viability in all phage preparations, suggesting the safety of LPRS-1 for potential applications in contaminated environments **(Fig 8D-F)**. This initial data serves as a foundational basis, implying that LPRS-1 could be a potent candidate for applications in various environments contaminated with MDR-*P. aeruginosa*.

**Figure 8:**
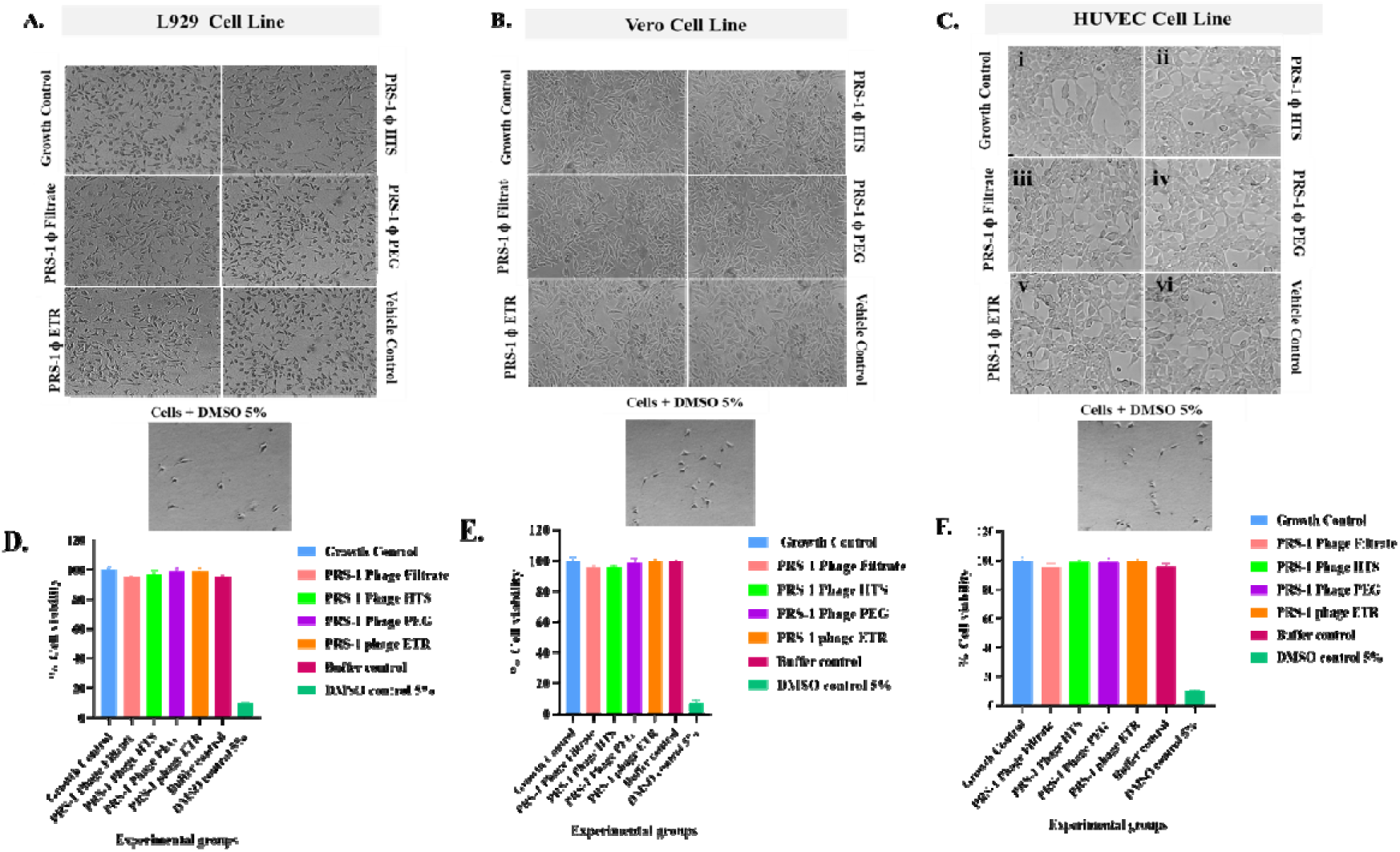
*In-vitro* cytotoxicity evaluation of phage PRS-1 preparations. **(A-C)** Inverted microscopic imaging of **(A)** L929 cells, **(B)** Vero cells, and **(C)** HUVEC cells treated with various preparations of phage and observed under 20 × objective. **(D-F)** Graphical representation showing quantification of the viable cells with MTT assay for the growth control (non-treated), Cytotoxic control (5% DMSO), and phagePRS-1 treated sets in **(D)** L929 cell lines **(E)** Vero cell lines **(F)** HUVEC cell lines.

## Conclusion

In summary, LPRS-1 emerges as a promising phage targeting MDR *P. aeruginosa*, including strain PAO1. It demonstrates potent antimicrobial activity against both planktonic cells and biofilms, achieving significant reductions in bacterial counts and effectively disrupting biofilm structures. Mechanistically, LPRS-1 modulates virulence factors of MDR *P. aeruginosa* by targeting QSS and carbapenemase expression, thereby addressing AMR. Importantly, =PRS-1 exhibits no *in-vitro* cytotoxic effects, indicating its potential safety in environmental applications. Furthermore, the application of =PRS-1 in reducing *P. aeruginosa* counts in raw sewage highlights the delivery of SDG 3, 6, 12, 14, 15 and its promise for future wastewater treatment strategies.

## Acknowledgments

Authors are thankful to the National Mission for Clean Ganga (NMCG), Government of India, New Delhi, India, for the project (GKC-01/2016-17, 212, NMCG-Research), Directors of CSIR-NCL, and CSIR-NEERI for infrastructure and support. RS acknowledges Mr. Manan Shah for his help with the sampling. RS is thankful to Mr. Mithil Mahale (Department of Chemistry, Savitribai Phule Pune University) and Mr. Abujunaid Khan (CSIR-NCL) for their guidance and help in the *in-vitro* cytotoxicity assays. RS is grateful to Mr. Tushar Kolhe and Mr. Chetan from Central Analytical Facility, CSIR-NCL, for their help in HR-TEM and FE-SEM. RS is thankful to HRDG-CSIR and NMCG, New Delhi, for their fellowship and AcSIR, New Delhi, for their academic support. RP acknowledges AcSIR for her academic support. The manuscript has been checked for plagiarism using iThenticate software, which has an institutional license.

## Declaration of Interest

The authors declare no conflict of interest. All authors have read and agreed to the submitted version of the manuscript.

## Credit authorship contribution statement

**RS:** Conceptualization, visualization, experimentation, sampling, writing-review & editing, and Data analysis. **SJ:** Isolation of MDR *P. aeruginosa* strains and experimentation. **RP:** qRT-PCR-related experimentation. **KK:** Project monitoring, Sample collection, editing, **MSD:** Conceptualization, Supervision, review, and editing

## Notes

### Competing Interest Statement

The authors have declared no competing interest.

